# The mitotic checkpoint protein MAD2 delivers monoamine transporters to endocytosis

**DOI:** 10.1101/2021.06.09.447721

**Authors:** Florian Koban, Michael Freissmuth

## Abstract

Monoamine transporters retrieve serotonin (SERT), dopamine (DAT) and norepinephrine (NET) from the synaptic cleft. Surface levels of transporters are also regulated by internalization. Clathrin-mediated endocytosis of cargo proteins requires adaptor protein 2 (AP2), which recruits cargo to the nascent clathrin-cage. The transporter C-terminus is required for internalization but lacks an AP2-binding site. In the present work, we show that internalization of SERT and DAT relies on MAD2, a protein of the mitotic spindle assembly checkpoint (SAC). A MAD2-interaction motif in the transporter C-terminus interacts with MAD2. This binding is contingent on the closed conformation of MAD2 and allowed for the recruitment of two additional SAC proteins, BubR1 and p31^COMET^ as well as AP2. MAD2, BubR1 and p31^COMET^ are present in serotoninergic neurons of the dorsal raphe, corroborating a biological role of the identified interactions. Depletion of MAD2 in HEK-293 cells stably expressing SERT and DAT decreases constitutive and triggered endocytosis, respectively. Altogether, our study describes a candidate mechanism, which connects monoamine transporters to the endocytic machinery and thus supports their internalization.

## INTRODUCTION

After their release into the synaptic cleft, the monoamine neurotransmitters serotonin, dopamine and norepinephrine are retrieved by their cognate transporters (serotonin transporter, SERT; dopamine transporter, DAT; norepinephrine transporter, NET), which reside in the presynaptic compartment in the vicinity of the active zone (Block *et al*, 2015; Hersch *et al*, 1997; Qian *et al*, 1995; Tao-Cheng & Zhou, 1999). By limiting diffusion, clearing the synapse and replenishing the vesicular pool of monoamine neurotransmitters, the transporters shape neurotransmission and neuromodulation. Their action translates in the regulation of e.g. mood, reward, movement, appetite or addictive behavior (Kristensen *et al*, 2011). SERT, NET and DAT are of conspicuous medical relevance, because they are the targets of approved (e.g. antidepressants, methylphenidate) and illicit drugs (e.g. cocaine, amphetamines). It has long been appreciated that the membrane-density of transporter molecules is also regulated by their endocytic removal from the cellular surface (Daniels & Amara, 1999; Melikian, 2004; Melikian & Buckley, 1999; Qian *et al*, 1997). The biological relevance of monoamine transporter internalization is not fully understood. However, transporter internalization can be triggered by exposure to their cognate substrate, amphetamines, antidepressants or the protein kinase C activator phorbol 12-myristate 13-acetate (PMA) (Daniels & Amara, 1999; Jayanthi *et al*, 2005; Jorgensen *et al*, 2014; Lau *et al*, 2008; Saunders *et al*, 2000). In addition, monoamine transporters also undergo constitutive internalization (Loder & Melikian, 2003; Rahbek-Clemmensen *et al*, 2014).

Simplified, two types of endocytosis can be distinguished, i.e. clathrin-dependent and -independent endocytosis. During clathrin-mediated endocytosis, a cage of clathrin triskelia locally assembles around the inner leaflet of the plasma membrane and stabilizes the formation of spherical pits, which ultimately pinch off as vesicles on the intracellular side. Both, triggered and constitutive internalization of DAT, depend on the action of clathrin (Sorkina *et al*, 2005). The molecular aspects of SERT endocytosis are less well explored, but it is reasonable to assume that internalization of these related transporters does not differ fundamentally. Clathrin-mediated endocytosis requires adaptor protein 2 (AP2), which links endocytic cargo to the clathrin coat. Hence, many cargos (e.g. the transferrin receptor) provide short amino acid motifs, which contact the adaptor protein. Classical adaptor protein binding motifs are either tyrosine (YXXϕ; X = any amino acid, ϕ = bulky hydrophobic residue) or dileucine ([DE]XXXL[LI]) based (Trowbridge *et al*, 1993). Some cargos rely on additional interacting proteins, which establish the contact to AP2 and other components of the endocytic machinery. The most prominent examples are the β-arrestins, which support the recruitment of AP2/clathrin to G protein-coupled receptors (Laporte *et al*, 1999; Oakley *et al*, 1999). The carboxyl terminus (C-terminus) of the dopamine transporter harbors a sequence (^587^FREKLAYAIA^596^), which is required for transporter endocytosis (Holton *et al*, 2005; Sorkina *et al.*, 2005). However, this sequence does not conform to a canonical AP2 interaction-site. This raises the possibility that DAT and related monoamine transporters rely on one or several auxiliary proteins, which support the interaction with AP2 and the clathrin coat in a manner reminiscent of arrestin-2 and -3.

During the cell cycle, a protein complex - referred to as the spindle assembly checkpoint (SAC) - safeguards correct chromosomal attachment to the mitotic spindle by sequestration of the anaphase promoting complex/cyclosome (APC/C) co-factor CDC20. Correct chromosome-alignment during metaphase causes disassembly of the SAC-complex, release of CDC20 and APC-mediated entry into anaphase. The small SAC protein MAD2 and its main interactors BubR1 and p31^COMET^ play a pivotal role in the SAC(Kops *et al*, 2020). Previous research found that, surprisingly, BubR1 interacted with β2-adaptin (a subunit of AP2) (Cayrol *et al*, 2002) and MAD2 with the C-terminus of the insulin receptor (O’Neill *et al*, 1997). More recently, MAD2 and BubR1 were shown to fulfill a moonlighting function as endocytic mediators by connecting the insulin receptor (IR) to the clathrin-coat (Choi *et al*, 2019; Choi *et al*, 2016): upon engagement of insulin, insulin receptor-bound MAD2 releases the inhibitory binding partner p31^COMET^ and recruits BubR1, which delivers the insulin receptor to AP2/clathrin-containing structures.

Interestingly, MAD2 is expressed in neuronal tissue of the human brain (O’Neill *et al.*, 1997; Uhlen *et al*, 2005; Uhlen *et al*, 2015; Yu *et al*, 2020). This raises the question on the role of a cell cycle protein in post-mitotic cells. In the present study, we show that monoamine transporters interact with the SAC-proteins MAD2, BubR1 and p31^COMET^. MAD2 binds directly to a MAD2-interaction motif in the C-terminus of the transporters. This binding is contingent on the “closed” conformation of MAD2. All three SAC-proteins are expressed in serotoninergic raphe neurons. Depletion of MAD2 in cells disrupts the interaction between SERT and BubR1/AP2 and decreases transporter endocytosis.

## RESULTS

### Neurotransmitter transporters contain putative C-terminal MIMs

The N- and C-termini of SLC6 transporters are more divergent than their hydrophobic cores; however, neurotransmitter transporters have conserved elements in their C-termini (Fig. EV1 A). MAD2-interaction motifs (MIMs) consist of a core motif of two hydrophobic residues, one basic residue (Arg or Lys) and a third hydrophobic residue followed by one or several prolines (Luo *et al*, 2002). Interestingly, C-termini of monoamine transporters harbor sequences which resemble MIMs (Fig. EV1 A, blue boxes). Furthermore, in DAT, this motif resides in a sequence previously shown to govern transporter-internalization (Holton *et al.*, 2005; Sorkina *et al.*, 2005) (Fig. EV1 A, green dashed box). Most of all, the SERT C-terminus contains a sequence element, which is consistent with a candidate MIM. We further corroborated this conjecture by aligning this sequence of SERT with MIMs of several canonical MAD2-binders (Fig. 1 A). In fact, the sequence in SERT was identical in all critical residues to the endogenous MAD2-binder ADAM17 (TACE) (Choi *et al.*, 2016; Nelson *et al*, 1999) and the unnatural peptide ligand MBP2 (MAD2 binding protein 2), which was discovered by phage display (Luo *et al.*, 2002). We therefore concluded that among neurotransmitter transporters, at the very least, the serotonin transporter might be a MAD2-binding protein. Thus, we first focused on SERT.

**Fig. 1.**
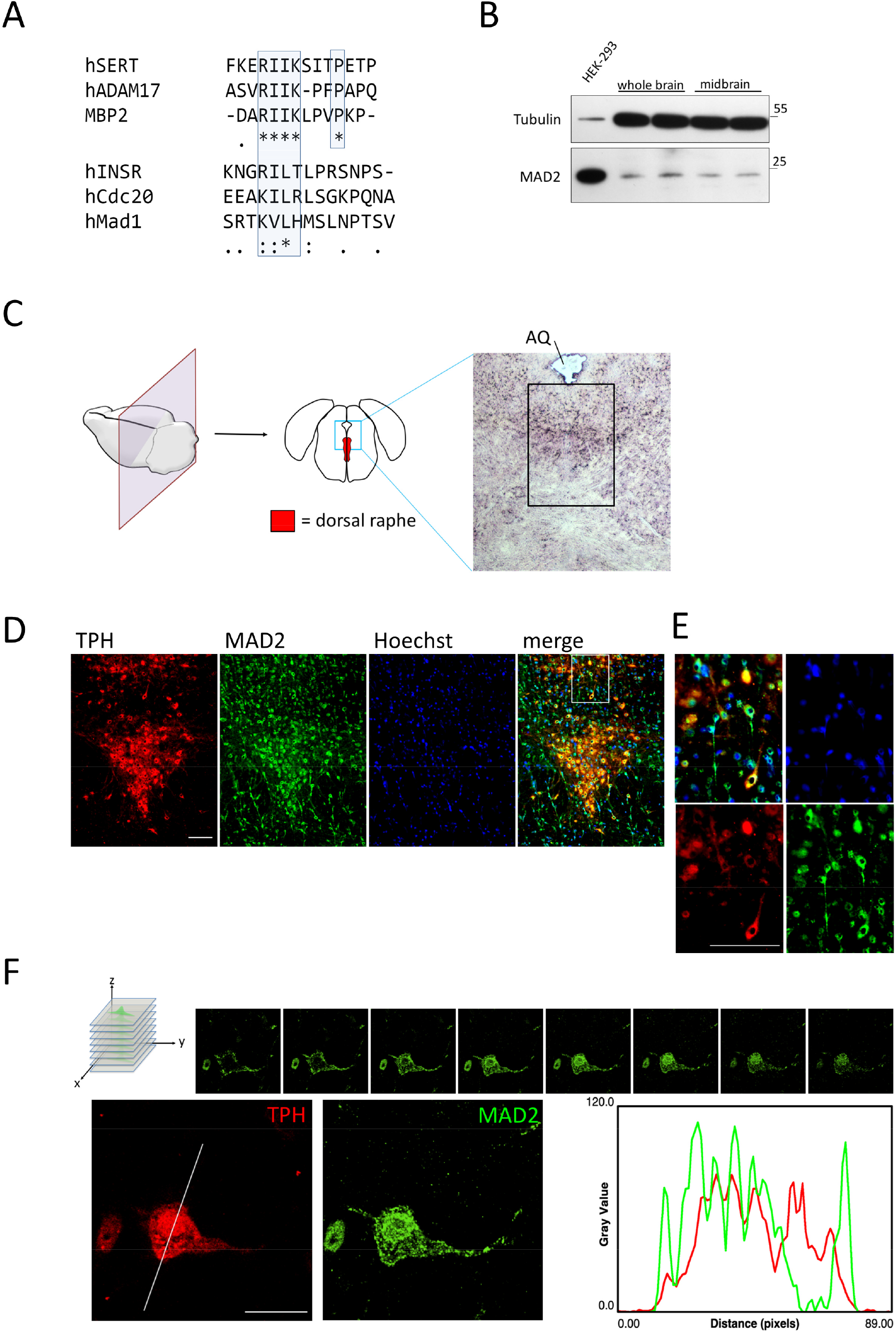
The serotonin transporter C-terminus harbors a potential MAD2 interaction motif (MIM). MAD2 protein expression in dorsal raphe neurons. A Clustal Omega alignment of the SERT C-terminus with previously described MAD2-interacting proteins. The putative SERT-MIM and previously described MIMs are highlighted by blue boxes; (**\***) - fully conserved residue; (:) - residues of strongly similar properties; (.) residues of weakly similar properties. B Whole brain and midbrain lysates of adult mice were prepared as outlined under “*Materials and Methods*”. Total protein (20 μg) was immunoblotted for MAD2 or α-Tubulin and compared to HEK-293 cell lysates. C Cryosections of mouse brain were prepared covering the dorsal raphe and stained with hematoxylin/eosin. The black box indicates the approximate area of immunofluorescence images. AQ = cerebral aqueduct. D – E Cryosections were subjected to immunofluorescence microscopy. The white box in the merge image represents the magnified area in (E). All images were taken as “multiple image alignments” using bright-field for H&E (10 × magnification) or appropriate excitation wavelengths and emission filters for immunofluorescence (20 × magnification). F Z-stacks of individual TPH+/MAD2+ positive neurons were generated by confocal microscopy. Intensity profiles of one representative optical plane were generated using ImageJ/Plot Profile software. Scale bars represent 100 μm in (D) and (E) and 20 μm in (F).

### MAD2-expression in mouse raphe neurons

It is a prerequisite that SERT and MAD2 are endogenously co-expressed in the same cell, if the putative interaction of SERT is of physiological relevance. First, we confirmed previous data (O’Neill *et al.*, 1997), that MAD2 can be found in whole brain, by immunoblotting lysates of mouse whole brain and midbrain. In fact, a band migrating at the same height as MAD2 from HEK-293 cells is present in all brain preparations (Fig. 1 B).

Accordingly, we examined serotoninergic neurons of the dorsal raphe nuclei for the presence of MAD2. We identified serotoninergic neurons by staining cryosections of mouse brain covering the dorsal raphe (Fig. 1 C) by double-immunofluorescence with antibodies against tryptophan hydroxylase (TPH) and MAD2 (Fig. 1 D). This approach verified the existence of MAD2 within TPH-positive dorsal raphe neurons. Magnification of a representative area (Fig. 1 E) showed that immunoreactive MAD2 was mainly cytosolic and was also visualized in neurite extensions. It is obvious that MAD2 was also found in TPH-negative cells throughout the whole extent of the section. This observation suggests that the protein fulfills tasks in the brain other than a putative interaction with the serotonin transporter. Control images, which demonstrate that the MAD2-signal is neither non-specific nor a bleed-through artifact of TPH in the respective cells, can be found as figure EV1 B.

The localization of MAD2 in TPH-positive neurons was explored by laser scanning confocal microscopy of individual TPH^+^/MAD2^+^ neurons along the Z-axis (Fig. 1 F): MAD2 was found in punctate structures in the cytosol and lining the cellular membrane. This distribution is consistent with the possibility that MAD2 interacts with neuronal membrane proteins.

### Binding of MAD2 to the candidate C-terminal MIMs

We verified an interaction of the C-terminus of SERT with MAD2 by employing fusion proteins of glutathione S-transferase (GST) harboring the C-terminus of SERT and of its closest relatives, DAT and NET (schematic representation of the constructs in Fig. 2 A). GST and GST fusion proteins immobilized on glutathione-conjugated beads were incubated with whole cell lysates prepared from HEK-293 cells. The immobilized material was analyzed for the presence of MAD2 by immunoblotting. After three washes, MAD2 was retained by GST fused to the SERT C-terminus (GST-SET-Ct) but not by GST alone (Fig. EV2 A, top pair of rows). When incubated with immobilized GST-DAT-Ct, the amount of retained MAD2 was substantially lower (Fig. EV2 A, bottom pair of rows), which can be accounted for by the fact that GST-DAT-Ct was prone to extensive degradation during purification (cf. Ponceau S stain in the bottom row of Fig. EV2 A). More importantly, during repetitive washes with detergent containing buffer, MAD2 was more readily released from GST-DAT-Ct than from GST-SERT-Ct (cf. first and third row in Fig. EV2 A). This suggests that MAD2 binds with higher affinity to the C-terminus of SERT than that of DAT.

**Fig. 2.**
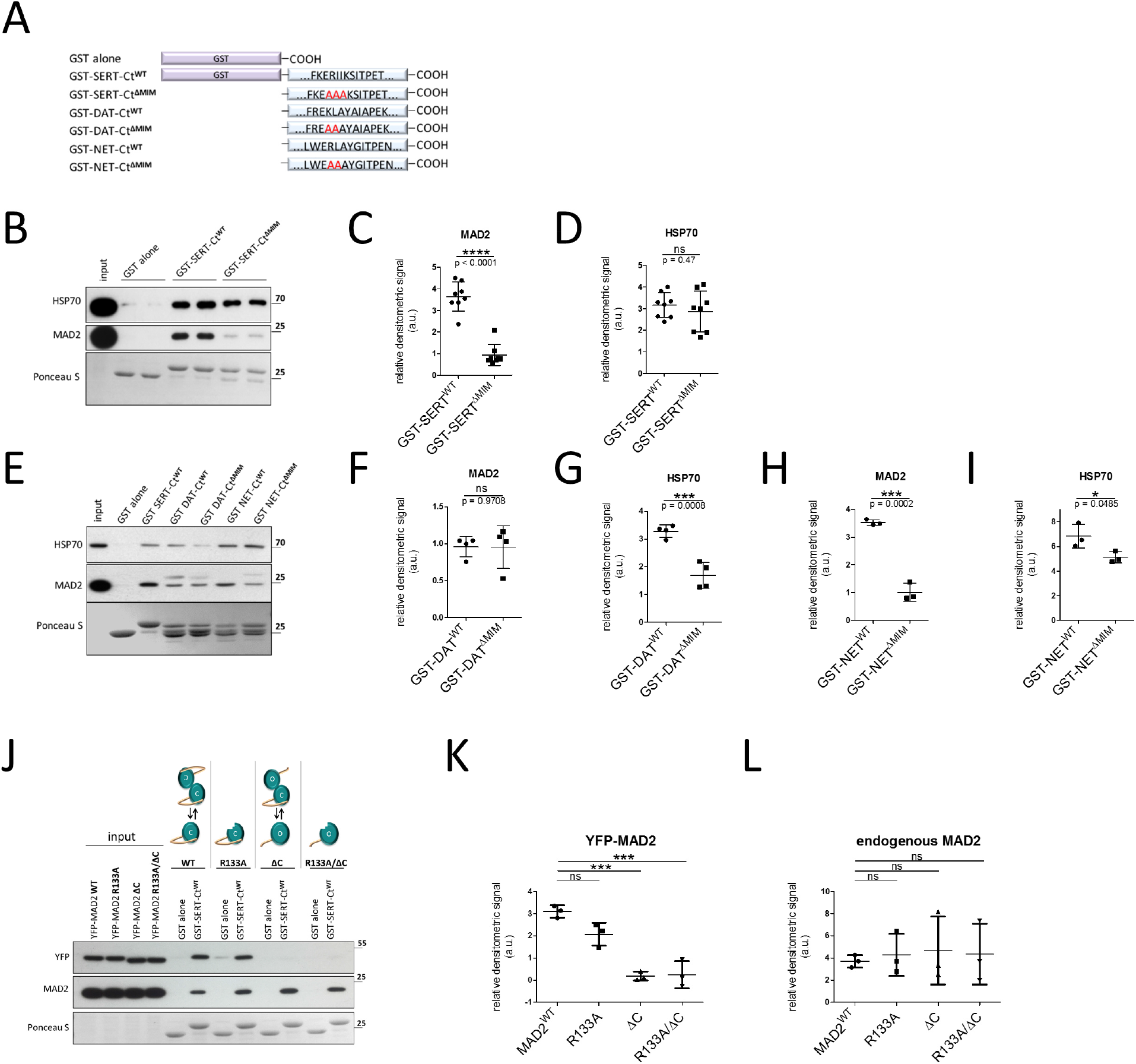
“Closed” MAD2 interacts with monoamine transporter C-termini at MAD2 interaction motifs. A Monoamine transporter C-termini fused to glutathione S-transferase (GST) and mutants thereof, which changed the putative MIM, were purified and used in GST-protein based interaction assays. B - I GST pull-down experiments were conducted in duplicates (B) or as single points (E) using the indicated GST fusion proteins. Eluates (30%) were subjected to immunoblotting together with an aliquot (1%) of the input. Blots were stained with Ponceau S and MAD2 was detected by immunoblotting. Contingent bands at around 30 kDa in (E) represent unspecific background signal deriving from the GST-protein. Densitometric signals were quantified using ImageJ software. Results were compared using unpaired two-tailed t-tests; *p < 0.05, ***p < 0.001, ****p < 0.0001, ns – not significant; exact p-values are indicated; error bars represent SD. J – L YFP-hMAD2-constructs were generated as outlined under “*Materials and Methods*”. Whole cell lysates of transfected HEK-293 cells were used for GST pull-down experiments. K – L One-way ANOVA with Dunnett’s multiple posthoc comparison; ***p < 0.001, ns – not significant; error bars represent SD.

We examined, if the bona fide MIM mediated the interaction, by replacing the Arg-Ile-Ile sequence with three alanine residues (RII-AAA) to generate GST-SERT-Ct^ΔMIM^ (Fig. 2 A, 3^rd^ construct). In fact, the mutation resulted in a significant drop in MAD2 binding (Fig. 2 B and C). HSP70 (heat shock protein of 70 kDA) binds to a segment in the SERT C-terminus preceding the MIM (El-Kasaby *et al*, 2014). Thus, as a control, we compared the ability of the wild type and mutated C-termini to retrieve HSP70: equivalent levels were observed (Fig. 2 C and D). This observation rules out that the mutation of the C-terminus compromised its ability to engage in protein-protein interactions by a non-specific effect.

We purified analogous GST fusion proteins for DAT and NET (Fig. 2 A, constructs 4 - 7). Interestingly, wild type GST-DAT-Ct and GST-DAT-Ct^ΔMIM^ bound equivalent levels of MAD2 (Fig. 2 E and F), but lower levels of HSP70 (Fig. 2 E and G). This suggests that, in the C-terminus of DAT, the HSP70 binding site extends into the putative MIM. In contrast, the C-terminus of NET bound MAD2 in a manner similar to that of SERT, i. e. the mutation in GST-NET-Ct^ΔMIM^ reduced MAD2 binding by ~70%, whereas HSP70 binding was significantly but only mildly affected (Fig. 2 E, H and I). Taken together, these observations confirm an interaction between the cell cycle protein MAD2 and the C-termini of monoamine transporters, which - at least for SERT and NET - is contingent on a canonical MIM.

### The closed conformation of MAD2 is the interacting state

MAD2 can adopt two thermodynamically stable conformations, referred to as open and closed (oMAD2 and cMAD2, respectively) (Mapelli *et al*, 2007; Yu, 2006). In the latter, the C-terminal region of MAD2 embraces the MAD2-interaction partner in a “safety belt”-like manner (Sironi *et al*, 2002). Previously described MAD2-interaction partners (e.g. CDC20, MAD1) and the insulin receptor form stable interactions with the closed conformation of MAD2 (Choi *et al.*, 2016; Luo & Yu, 2008). It is reasonable to surmise that the interaction between SERT and MAD2 is governed by the same principle. We verified this prediction by generating YFP-tagged MAD2 and by incorporating a series of mutations. The YFP-tag was chosen for two reasons: (i) it adds about 25 kDa to the size of MAD2 (~ 23 kDa). Hence, the resulting tagged protein can readily by distinguished from endogenous MAD2 on the same immunoblot (Fig. EV2 B). (ii) The commercially available MAD2-antibody is directed against the C-terminus and does not recognize a C-terminal truncation mutant of MAD2 (Fig. EV2 B). MAD2 requires an intact C-terminus to adopt the closed conformation. This strict requirement allows for the generation of a constitutively open MAD2 by deleting the last 10 amino acids (MAD2^ΔC^) (Luo *et al*, 2000).

Closed MAD2 dimerizes with both, another cMAD2 (symmetric dimer) and oMAD2 (asymmetric dimer). In contrast, ligand-bound cMAD2 can only form asymmetric dimers (Luo & Yu, 2008; Yang *et al*, 2008). Mutation of arginine at position 133 to alanine (MAD2^R133A^) renders MAD2 dimerization-deficient (Sironi *et al*, 2001). This is useful to investigate, if binding of MAD2 interaction partners depends on MAD2 dimerization. MAD2^ΔC/R133A^ was also generated to rule out that open MAD2^ΔC^ indirectly associates with the transporter C-terminus merely by dimerization with ligand-bound endogenous cMAD2.

YFP-tagged versions of MAD2 were transiently expressed in HEK-293 cells and lysates thereof were subjected to GST pull-down experiments as described above. As evident from figures 2 J and K, equivalent levels of wild type YFP-MAD2 and of monomeric YFP-MAD2^R133A^ were retrieved by GST-SERT-Ct. This indicates that the dimeric nature of MAD2 is immaterial for binding to SERT. In contrast, both YFP-MAD2 constructs lacking the C-terminus failed to bind to GST-SERT-Ct (Fig. 2 J, K). Thus, the closed conformation of MAD2 is required to support its interaction with SERT. In addition, the observations indicate that MAD2 binds to SERT as a monomer, because MAD2^ΔC^ was not pulled down concomitantly with endogenous MAD2 (cf. top and middle row in Fig. 2 J). Finally, we stress that the pull-down of endogenous MAD2 allowed for an internal control: under all conditions, similar levels of endogenous MAD2 were retrieved by GST-SERT-Ct (Fig. 2 J and L).

### BubR1 and p31^COMET^ are expressed in raphe neurons and interact with transporter C-termini

As part of the spindle assembly checkpoint, MAD2 interacts with numerous proteins. When bound to the insulin receptor, MAD2 recruits the SAC proteins BubR1 and p31^COMET^ (Choi *et al.*, 2016). We examined, if this was also the case for the C-termini of the monoamine transporters: in GST pull-down assays, BubR1 and p31^COMET^ were retrieved by the wild type C-terminus of SERT, DAT and NET (Fig. EV3 A). Disruption of the MIM of SERT and NET (ΔMIM-containing constructs) resulted in a pronounced reduction in immunoreactive BubR1 and p31^COMET^; this effect was less apparent for DAT.

We verified the expression of BubR1 and p31^COMET^ in dorsal raphe neurons. Both proteins were detected by immunofluorescence microscopy using specific antibodies in TPH+ serotoninergic raphe neurons. In addition - and similar to MAD2 - BubR1 and p31^COMET^ were not confined to TPH+ neurons: they were also seen in the area adjacent to the raphe nucleus (Fig. 3 A and C). Images captured at higher magnifications showed that immunoreactivity for BubR1 was distributed uniformly in the cytosol of TPH^+^ neurons. We rule out that the cytosolic BubR1 signal is a bleed-through artifact deriving from the TPH-signal: cytosolic BubR1 was readily detectable in cells, which were devoid of any immunoreactivity for TPH (Fig 3 B, white arrowheads). In contrast to BubR1, immunostaining for p31^COMET^ was enriched in a region surrounding the nucleus of TPH^+^ neurons, but it was also detectable in the cytosol (Fig. 3 C and D).

**Fig. 3.**
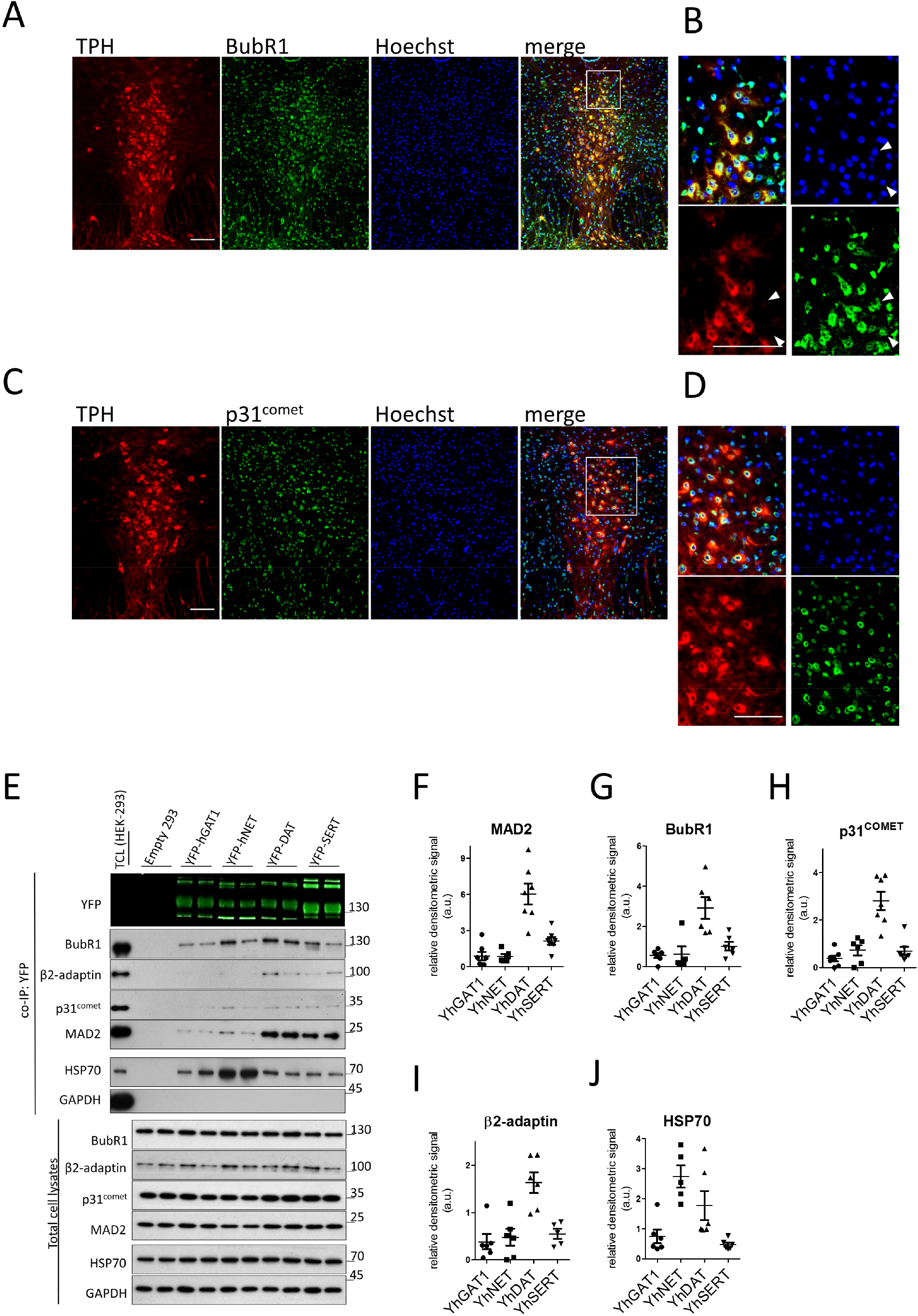
Monoamine transporters associate with β2-adaptin and the SAC-proteins p31^COMET^ and BubR1, which are expressed in raphe nuclei. A – D Immunofluorescence of p31^COMET^ and BubR1 in mouse dorsal raphe. White boxes in the merge images represent the magnified area in (B) and (D). White arrowheads in (B) point to cytosolic BubR1 in TPH-negative cells. Scale bars represent 100 μm. E – J Co-immunoprecipitation experiments using lysates from HEK-293 cells stably expressing the indicated YFP-tagged transporters. Eluates (20%) as well as 0.1% total cell lysate were analyzed by immunoblotting. The immunoblot for mouse anti-β2-adaptin was subsequently used for incubation with rabbit anti-GFP primary and IRDye 800CW anti-rabbit secondary antibody. IRDye signals were detected using the LI-COR Odyssey CLx imaging system. Densitometric signals were quantified using ImageJ software. All signals were normalized to their respective transporter signal intensities; error bars represent SD.

### Correlation between co-immunoprecipitated SAC-proteins and clathrin adaptors

We immunoprecipitated heterologously expressed YFP-tagged variants of SERT, DAT, NET and GAT1 to verify, if full-length neurotransmitter transporters formed complexes with SAC-proteins in living cells. Because of the similar molecular mass, the light chain of IgG used for conventional co-IPs may interfere with the immunodetection of MAD2 (~23 kDa). Accordingly, we resorted to anti-GFP nanobodies covalently conjugated to agarose (GFP-Trap) for retrieving YFP-tagged transporters from detergent lysates of stably transfected HEK-293 cells. MAD2 was co-immunoprecipitated with all transporters, albeit at very different amounts (Fig. 3 E and F). Similarly, variable levels of BubR1 and p31^COMET^ were found in the immunoprecipitates (Fig. 3 E, G and H). We used complex formation between HSP70 and newly synthesized transporters as a positive control (Asjad *et al*, 2017; El-Kasaby *et al*, 2019; El-Kasaby *et al.*, 2014; Kasture *et al*, 2016): HSP70 was co-immunoprecipitated by each transporter (Fig. 3 E, J). As a negative control, we used the abundant cytosolic protein glyceraldehyde 3-phosphate dehydrogenase (GAPDH): this protein was not detected in the immunoprecipitates (Fig. 3 E). Hence, we conclude that the observed co-immunoprecipitation of MAD2, BubR1 and p31^COMET^ with the transporters reflected a specific association.

If MAD2 drives the interaction of the spindle assembly checkpoint (SAC) proteins BubR1 and p31^COMET^, their levels in the co-immunoprecipitate are predicted to be correlated. This was the case: we observed a statistically significant correlation between the amount of co-immunoprecipitated p31^COMET^ and BubR1 and the amount of transporter-associated MAD2 (Fig. EV3 B and C). Furthermore, a correlation was also seen for the co-immunoprecipitated amount of the AP2-subunit β2-adaptin and MAD2 (Fig. 3 E and Fig. EV3 D). In contrast, the levels of co-immunoprecipitated HSP70 did not show any correlation with the amount of MAD2 (Fig. EV3 E). Taken together, these observations confirm that neurotransmitter transporters - in particular DAT and SERT- interact with proteins of the spindle assembly checkpoint. The correlation between co-immunoprecipitated levels of BubR1, p31^COMET^, AP2 and of MAD2 suggests a complex between SAC-proteins and proteins of the clathrin endocytic machinery, which may drive endocytosis of neurotransmitter transporters.

### MAD2 mediates SERT/SAC/AP2 complex formation

If monoamine transporters internalized via a MAD2-mediated module of SAC-proteins and endocytic proteins, elimination of MAD2 ought to cause disassembly of this complex. The GST pull-down experiments provided circumstantial evidence to support this conjecture, because they showed that the interaction with p31^COMET^ and BubR1 also required the MAD2 interaction motif in the C-terminus of SERT and NET (Fig. EV3 A). We obtained evidence that is more direct by depleting MAD2 in HEK-293 stably expressing YFP-SERT using siRNA-mediated knockdown followed by co-immunoprecipitation (Fig 4 A). Knockdown of MAD2 resulted in a substantial reduction of BubR1 co-immunoprecipitated with SERT (Fig. 4 B) and a drop by about 50% of β2-adaptin (Fig. 4 C). The levels of p31^COMET^ were tendentially reduced but the difference did not reach statistical significance (Fig. 4D). As an alternative approach, we examined the consequences of MAD2 knockdown for binding of p31^COMET^ in a GST pull-down experiment (Fig 4 F): siRNA-mediated depletion of MAD2 from the total cell lysate *per se* reduced the amount of p31^COMET^ retained by the C-terminus of SERT. This effect was further enhanced by the mutation of the MAD2-interaction motif: negligible levels of p31^COMET^ were retrieved from MAD2-depleted lysates by GST-SERT-Ct^ΔMIM^ (Fig. 4 F and G). Hence, we conclude that MAD2 supports the recruitment of BubR1, p31^COMET^ and AP2 to the serotonin transporter.

**Fig. 4.**
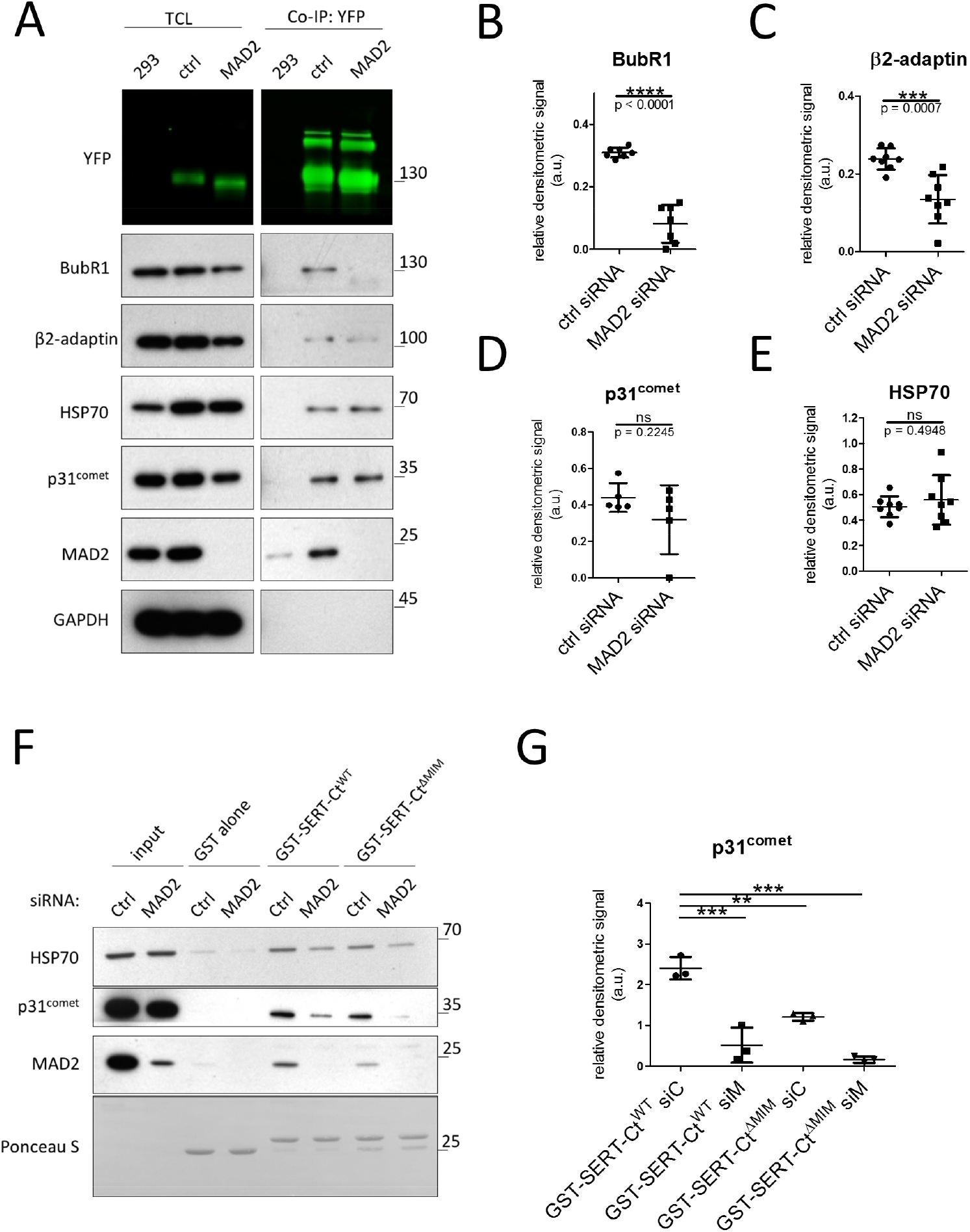
MAD2-depletion causes SERT/SAC/AP2-complex disassembly. A – E HEK-293 cells stably expressing YFP-hSERT were transfected with indicated siRNAs and subjected to co-immunoprecipitation. Results were compared using unpaired two-tailed t-tests; ***p < 0.001, ****p < 0.0001, ns – not significant; exact p-values are indicated; error bars represent SD. F – G HEK-293 cells were transfected with the indicated siRNAs and cell lysates were used for GST pull-down experiments using the indicated GST-proteins. One-way ANOVA with Dunnett’s multiple posthoc comparison; **p < 0.01, ***p < 0.001; error bars represent SD.

### MAD2 depletion increases SERT surface expression and blocks intracellular SERT accumulation

Based on the data summarized in figures 1 – 4, we inferred a role of MAD2 in driving clathrin-mediated endocytosis of monoamine transporters. The velocity of substrate uptake correlates with surface presence of the transporters. Hence, substrate uptake is predicted to be enhanced, if MAD2-depletion precludes constitutive endocytosis and thus results in the accumulation of SERT at the cell surface. Accordingly, we transfected HEK-293 cells stably expressing YFP-SERT with siRNAs against MAD2 and measured serotonin uptake. As shown in figure EV4 A, MAD2 knockdown was efficient. The K_M_ of the substrate reflects the binding of serotonin and co-substrate ions and is determined by all subsequent rates of conformational transitions, which afford the cytosolic release of substrate and sodium and return of the transporter to the outward facing state (Burtscher *et al*, 2019). MAD2 depletion did not alter the K_M_ of serotonin transport (Fig. EV4 B). Hence, we consider it unlikely that MAD2 impinges on the rates, which govern the transport cycle. However, MAD2 depletion increased V_max_ by about 40%. An increased density of SERT at the plasma membrane can account for this enhanced maximum rate of transport. Accordingly, we examined the total amount of SERT in detergent lysates and its surface levels in control cells and in cells, where MAD2 had been depleted: immunoblotting revealed, that after siRNA-mediated knockdown of MAD2, SERT levels were elevated in both, detergent lysates and in the SERT pool, which was at the cell surface and thus accessible to the membrane-impermeable biotinylation reagent (Fig. EV4 C). Visual inspection by confocal microscopy revealed intracellular accumulation of SERT in control cells. This intracellular pool of SERT was seemingly absent in MAD2-depleted cells (Fig. 5 A). Quantification confirmed that intracellular SERT was present in significantly lower amounts in cells transfected with MAD2 siRNA (Fig. 5 B).

**Fig. 5.**
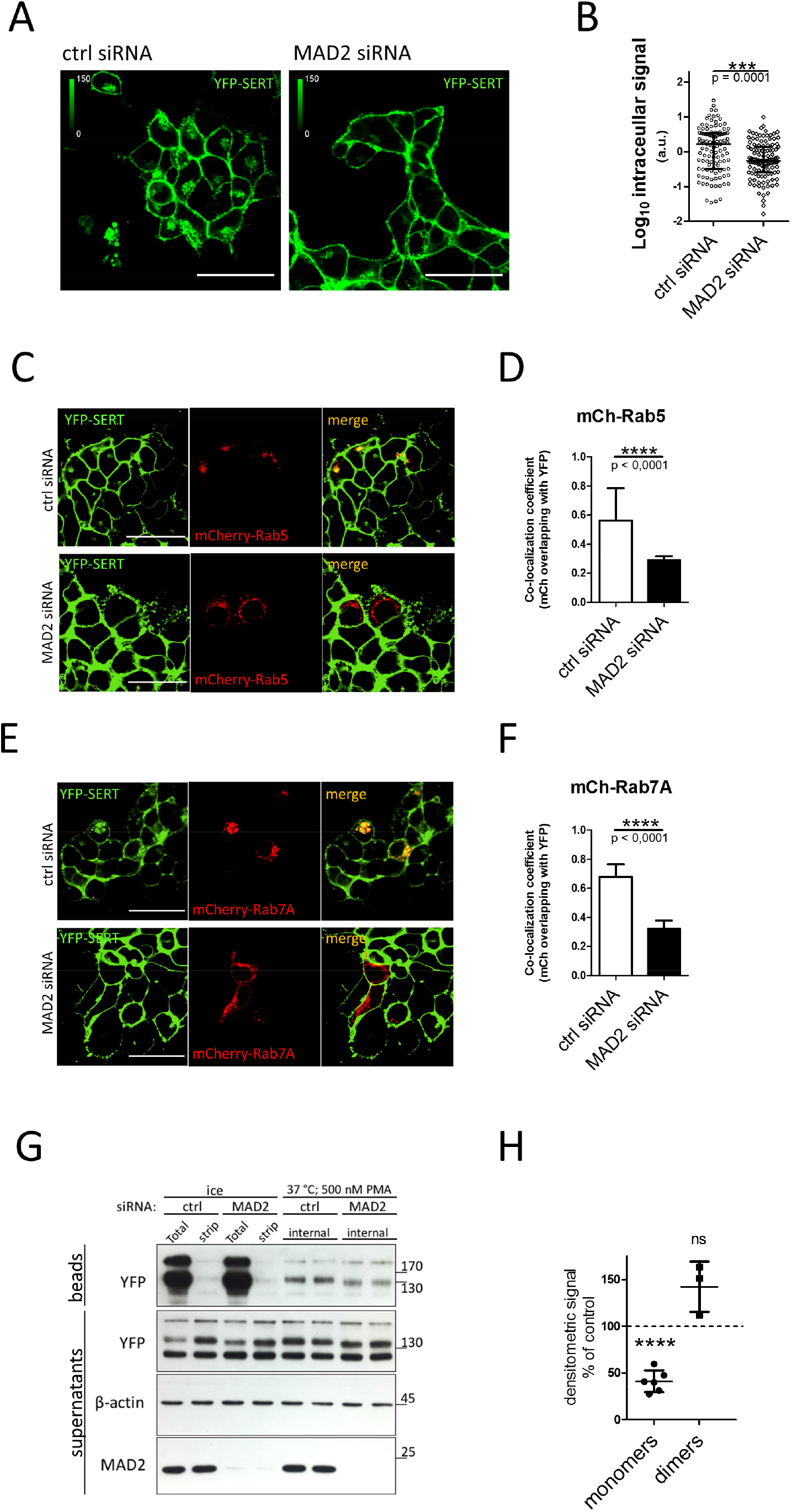
MAD2-depletion enriches SERT at the cell surface and impairs transporter endocytosis. A - B HEK-293 cells stably expressing YFP-hSERT were transfected with the indicated siRNAs and imaged and analyzed as described in “*Materials and Methods*”. Ctrl siRNA = 112 cells; MAD2 siRNA = 116 cells. Values are presented as Log_10_ to account for the large variation in the signal. Unpaired two-tailed t-test, ***p < 0.001; exact p-value is indicated. Shown is the median and the interquartile range. Scale bars represent 50 μm. C – F HEK-293 cells stably expressing YFP-hSERT were siRNA-transfected, co-transfected with plasmids encoding mCherry-Rab5 or mCherry-Rab7A and imaged as described in „Materials and Methods“. Manders’ correlation coefficients for the fraction of mCherry overlapping YFP were calculated for multiple individual images using ImageJ/JACoP. Number of analyzed images: mCh-Rab5/ctrl-siRNA = 38; mCh-Rab5/MAD2-siRNA = 33; mCh-Rab7A/ctrl-siRNA = 13; mCh-Rab7A/Mad2-siRNA = 14. Unpaired two-tailed t-test, ****p < 0.0001. Error bars represent SD. Scale bars represent 50 μm. G – H HEK-293 cells stably expressing YFP-hDAT were used for reversible biotinylation experiments. “Internalization” samples were prepared in duplicates, “total” and “strip” samples as single points. Eluates (30%) and supernatants (1%) were analyzed by standard immunoblotting. Densitometric signals for YFP-DAT dimers are inherently weaker than for YFP-DAT monomers. Hence, only half of the experiments produced results, which allowed for an analysis of the dimeric species. Results for monomeric and dimeric YFP-hDAT signals are expressed as percent of “ctrl siRNA” (dashed line) and compared using paired two-tailed t-tests, ****p < 0.0001, ns – not significant; error bars represent SD.

### MAD2 is required for transporter internalization

SERT is constitutively internalized and sorted to the late endosomal and lysosomal compartment (Rahbek-Clemmensen *et al.*, 2014). Hence, intracellular SERT-positive structures may be - at least in part - of endocytic origin. We identified these structures by transfecting YFP-SERT expressing cells with plasmids encoding mCherry-Rab5 (an early endosomal marker) and mCherry-Rab7A (a late endosomal marker). In HEK-293 cells subjected to transfection with control siRNA, intracellular SERT clearly co-localized with both endosomal markers (upper panels in Fig. 5 C and E). This was not observed in cells transfected with MAD2 siRNA (lower panels in Fig. 5 C and E). Quantification of co-localization over multiple individual images confirmed that YFP-SERT overlaps with Rab5 or Rab7A to significantly higher extents in MAD2-depleted cells.

We verified that depletion of MAD2 impaired transporter internalization by reversibly biotinylating surface proteins. The experimental strategy is outlined by the schematic representation in Fig. EV4 D: after labeling of surface protein with the biotinylation reagent, internalized proteins are protected from subsequent cleavage of the biotin moiety by TCEP (tris(2-carboxyethyl)phosphin). The protein kinase C activator phorbol 12-myristate 13-acetate (PMA) triggers clathrin-mediated endocytosis of SERT (Qian *et al.*, 1997) and DAT (Daniels & Amara, 1999; Melikian & Buckley, 1999). However, in HEK-293 cells, the effect is more robust with DAT, i.e. a larger fraction of DAT is internalized than of SERT (cf. refs. 7 and 8) and this internalization occurs within 10 minutes (Gabriel *et al*, 2009). Hence, internalization was triggered by exposing HEK-293 cells stably expressing YFP-tagged DAT to 500 nM PMA at 37°C. Consistent with previous reports, we observed that PMA resulted in the accumulation of a cleavage-resistant, biotinylated fraction of DAT (duplicate samples in lanes 5 and 6 of Fig. 5 G). This fraction was significantly reduced in cells, in which MAD2 was depleted by siRNA-mediated knockdown (duplicate samples in lanes 7 and 8 of Fig 5 G; Fig. 5 H). Thus, a reduction in cellular MAD2 levels blunted PMA-triggered internalization of DAT. This observation is consistent with a role of MAD2 during clathrin-mediated endocytosis. Interestingly, only endocytosis of the transporter species, which migrated as monomer (at ~110 kDa), was impaired by MAD2 depletion. In contrast, comparable levels of the species migrating above 170 kDa - corresponding to apparent dimers - were internalized in control and MAD2-depleted cells (Fig 5 H). These results are in line with recent data, which show that dopamine transporters driven to oligomerize via the action of the small molecule AIM-100 undergo endocytosis by a clathrin-independent mechanism (Sorkina *et al*, 2018).

## DISCUSSION

The generalized model of clathrin-mediated endocytosis posits that adaptor protein 2 (AP2) recognizes short sequence motifs on client molecules. These allow for recruitment of client proteins to clathrin-coated membrane areas and for their subsequent internalization. An additional adapter is required, if these sequence motifs are not present. The C-terminus of the dopamine transporter contains a sequence element, which is indispensable for clathrin-mediated internalization (Holton *et al.*, 2005; Sorkina *et al.*, 2005). However, the motif does not qualify for direct AP2-binding. Hence, the molecular mechanism underlying clathrin-mediated endocytosis of DAT has remained elusive. This is also true for other neurotransmitter transporters (e.g. SERT, NET, GAT1), which are known to undergo internalization. In the present work, we identified MAD2 as a crucial component required to drive clathrin-mediated endocytosis of SERT and DAT. This conclusion is based on the following lines of evidence: (i) our experiments show that the C-termini of monoamine transporters harbor an interaction site for MAD2, which allows for binding of MAD2 and its interaction partners, BubR1 and p31^COMET^. (ii) Complex formation between transporters and these three spindle assembly checkpoint proteins was confirmed by co-immunoprecipitation and was shown to be contingent on the presence of MAD2. These complexes also contained the AP2 component β2-adaptin and thus provide the link to the clathrin coat. (iii) Constitutive internalization of SERT and regulated endocytosis of DAT required the presence of MAD2. (iv) The spindle assembly checkpoint proteins were visualized in serotoninergic neurons. To the best of our knowledge, the current findings are the first to provide a mechanism, which affords the communication of transporter C-termini with the components of the endocytic machinery.

Our observations also shed light on the question, why spindle assembly checkpoint proteins should be present in post-mitotic neurons of the brain. We suspect that MAD2 has additional client proteins, which are trafficked as endocytic cargo. MAD2 was not only visualized in the cytosol (or in association with the nucleus) but also in punctate structures along the membrane of the soma and neuronal extensions. This localization is in agreement with a cellular function of MAD2 during membrane trafficking. Some two decades ago, the mRNA of MAD2 was found to be more abundant in human brain (and other non-dividing tissues such as skeletal muscle and the heart) than in tissues, where cell division was expected to occur (e.g. lung and liver) (O’Neill *et al.*, 1997). Furthermore, the “human protein atlas”-project also revealed that, in human brain tissue, MAD2 was present in neurons but not in glial cells; the subcellular distribution was also described as cytoplasmic and membranous (https://www.proteinatlas.org/ENSG00000164109-MAD2L1/tissue/cerebral+;cortex#img) (Uhlen *et al.*, 2005; Uhlen *et al.*, 2015). More recently, expression analysis of several mouse tissues revealed that virtually all SAC-proteins (including MAD2, BubR1 and p31^comet^) are expressed in terminally differentiated neurons and adult brain (Yu *et al.*, 2020). It is worth noting that BubR1 and its relative BUB1 also bind adaptins other than β2-adaptin (Cayrol *et al.*, 2002). Hence, we propose that MAD2 and BubR1/BUB1 play a more widespread role in vesicular sorting than merely in endocytosis of neurotransmitters of the SLC6 family and of the insulin receptor.

Binding of MAD2 to the C-terminus of SERT required its closed conformation. This observation is consistent with the conformational requirement for binding of MAD2 to the insulin receptor (Choi *et al.*, 2016). MIMs adopt a β-strand structure, which allows for some variation in their sequence (Luo *et al.*, 2002). However, in the crystal structures of SERT and DAT, the region comprising the MAD2-interaction motif lies within an amphipathic α-helix, which is stabilized by a cation-π interaction and a salt bridge with intracellular loop 1 in DAT and SERT, respectively (Coleman *et al*, 2016; Koban *et al*, 2015; Penmatsa *et al*, 2013). It has been proposed that phosphorylation (and/or other modifications) may change the conformation of the C-terminus and thus allow for access of interacting proteins (Coleman *et al.*, 2016; Koban *et al.*, 2015; Penmatsa *et al.*, 2013). Accordingly, the conformational flexibility of the C-terminus of DAT and SERT provides a means for a regulated association of MAD2, which initiates clathrin-mediated endocytosis of the transporter.

SERT (and all related neurotransmitter transporters) exert their eponymous action, i.e. the retrieval of released neurotransmitter, in the presynaptic specialization. The fact that we visualized MAD2 in neurite extensions of serotoninergic neurons suggests that MAD2 can also reach axonal endings and thus support clathrin-mediated endocytosis in synaptic boutons. However, SERT is also delivered to the surface of the neuronal soma. Somatodendritic SERT also supports serotoninergic signaling (Colgan *et al*, 2012; Kasture *et al*, 2019). This is also true for somatodendritic DAT (Falkenburger *et al*, 2001). Thus, while a role of MAD2 during endocytosis of axonal SERT and DAT remains to be demonstrated, the currently available evidence supports the conclusion that MAD2-dependent regulation of SERT and DAT in the neuronal soma is of biological relevance.

## MATERIALS AND METHODS

### Protein sequence alignments

Protein sequences were aligned using Clustal Omega software (Sievers *et al*, 2011).

### Cell culture

Generation of HEK-293 cells stably expressing N-terminally YFP-tagged human isoforms of SERT (YFP-hSERT), DAT (YFP-hDAT) and NET (YFP-hNET) was described previously (Mayer *et al*, 2016; Niello *et al*, 2019). HEK-293 cells stably expressing YFP-hGAT1 were generated accordingly. In brief, cells were transfected with YFP-hGAT1 using jetPRIME (114-15, Polyplus-Transfection). Stably expressing cells were selected in the presence of geneticin (250 μg mL^−1^) and enriched using fluorescence-activated cell sorting. All cell lines were maintained in humidified atmosphere at 37°C 5% CO_2_ in antibiotic-free high glucose Dulbecco′s modified Eagle′s medium supplemented with 10% fetal bovine serum. Generation of the plasmid encoding YFP-hGAT1 is described in section “Plasmids, Molecular Cloning, Mutagenesis”.

### Antibodies

#### Immunofluorescence

<u>Primary antibodies</u>: rabbit-anti-MAD2 (1:200; ab70385, abcam); sheep-anti-tryptophan hydroxylase/TPH (1:150; ab32821, abcam); rabbit-anti-BubR1 (1:100; A300-386A, Bethyl Laboratories Inc.); rabbit-anti-P31^COMET^ (murine) (1:100; provided by Dr. Hongtao Yu; UT Southwestern, Dallas, Texas, USA). <u>Secondary antibodies</u>: Alexa Fluor 488 donkey-anti-rabbit IgG (1:500; A21206, Invitrogen); Alexa Fluor 555 donkey-anti-sheep IgG (1:500; A21436, Invitrogen).

#### Immunoblotting

<u>Primary antibodies</u>: rabbit-anti-MAD2 (1:2500; ab70385, abcam); mouse-anti-αTubulin DM1A (1:2000, T9026-100UL, Sigma-Aldrich); mouse-anti-HSP70 (1:3000; ab47455, abcam); rabbit-anti-GFP (1:5000; ab290, abcam); rabbit-anti-BubR1 (1:1500; A300-386A, Bethyl Laboratories Inc.); mouse-anti-βActin (1:2000; A0760-40, USBiological Life Sciences); mouse-anti-Adaptin β (1:500; 610382, BDbiosciences); mouse-anti-GAPDH 0411 (1:2000; sc-47724, SantaCruz Biotechnology); rabbit-anti-p31^COMET^ (human) (1:1000; provided by Dr. Hongtao Yu; UT Southwestern, Dallas, Texas, USA). <u>Secondary antibodies</u>: HRP-linked anti-mouse IgG (1:5000; 7076S, Cell Signaling); HRP-linked anti-rabbit IgG (1:5000; 7074S, Cell Signaling); IRDye 800CW goat-anti-Rabbit IgG (1:5000; 925-32211, Li-Cor).

### Tissue microscopy

Male C57BL/6 mice (10 weeks of age) were killed by decapitation under deep anesthesia with isoflurane. Brains were excised, covered with Tissue-Plus O.C.T compound (4583, Scigen), snap-frozen in liquid N_2_ and stored at −80°C. Cryosections (10 μM), covering the dorsal raphe nuclei, were prepared on a cryostat, immediately subjected to immunostaining or frozen at −80°C. Sliced tissue was fixed in acetone/methanol (1:1) for 15 min at −20 °C and embedding matrix was removed in ddH_2_O. Slides were attached to Shandon coverplates (Thermo Scientific) and inserted into a Shandon Sequenza holder. After washing in TBS (20 mM Tris-HCl, pH = 7.6; 150 mM NaCl), tissue was blocked in 5% normal goat serum + 5% normal donkey serum for 1 h at room temperature. Subsequently, tissue was incubated with indicated primary antibodies (as outlined under “Antibodies”) at 4 °C overnight. After 3 wash steps in TBS, sections were incubated in fluorescent secondary antibodies for 1 h at room temperature, together with Hoechst 33342 (14533, Sigma Aldrich) at 1:3000 for nuclear staining. All blocking and antibody dilutions were prepared in Dako antibody diluent (Agilent). After three wash steps in TBS, slides were mounted in Fluoromount-G Mounting Medium (00-4958-02, Invitrogen) and covered with glass cover slips. Fluoromount-G solidified overnight under brass weights (~100 g per slide).

Alternatively, tissue sections were stained with hematoxylin/eosin (H&E) using a standard protocol (publically available from the Adler lab, Johns Hopkins University).

Mounted tissue sections were imaged on an Olympus AX70 epifluorescence microscope using a magnification of 10- and 20-fold for H&E stainings and for immunofluorescence, respectively. Adjacent single images were acquired in order to afford multiple image alignments (MIA) arranged by Olympus Cell^P software. In addition, identical slides were imaged on a Zeiss LSM 510 laser scanning confocal microscope equipped with Plan-Apochromat 63×/1.4 oil-immersion objective. Either single optical planes were acquired or Z-stacks with a slice thickness of 0.4 μM. Images were analyzed using ImageJ/Plot Profile software.

### Immunoblotting of mouse brain proteins

Male C57BL/6 mice (23 weeks of age) were killed by decapitation under deep anesthesia with isoflurane and their brains were excised. For immunoblots of midbrain proteins the respective area was separated from the residual brain. Tissues were homogenized in 2 mL co-IP lysis buffer (20 mM Tris-HCl, pH = 7.6; 150 mM NaCl; 1 mM EDTA; 10% glycerol; 1% Nonidet P40 Substitute; protease inhibitor cocktail [Roche]; PhosSTOP [Roche]), using a Dounce homogenizer. Lysates were cleared at 16 100 g for 30 minutes at 4°C. Protein concentrations were measured by dye binding (Coomassie Brilliant Blue G-250) and diluted to 1 mg mL^−1^ in co-IP lysis buffer. Samples (20 μg) were analyzed by standard SDS-PAGE and immunoblotting.

### GST-Protein Purification and GST Pull-down

The constructs for the GST-tagged C-terminus of human NET and human DAT were generated as described in section “Plasmids, Molecular Cloning, Mutagenesis”. GST-proteins were purified from transformed XL10-Gold *Escherichia Coli* as described previously (El-Kasaby *et al.*, 2014). For GST-protein based interaction assays, 25 μg GST-proteins were pre-bound to glutathione sepharose (20 μL packed resin per sample) in 200 μL TBS-T for 2 h at room temperature under end-over-end rotation. Meanwhile, adherent HEK-293 cells on confluent 10 cm dishes (approx. 9 × 10^6^ cells per dish) were lysed in 450 μL ice-cold co-IP lysis buffer. Lysates were cleared at 16 100 g at 4°C for 30 min. The supernatant was recovered and used for GST pull-down assays.

Immobilized GST-proteins were washed once in 500 μL co-IP lysis buffer and incubated with cleared HEK-293 lysate (150 μL ≙ ~ 3 × 10^6^ cells per sample) for 1 h at room temperature. Associated proteins were washed 3 times in 500 μL ice-cold TBS-T and eluted in 50 μL conventional 2× Laemmli SDS-PAGE sample buffer at 90°C for 10 min. In the same step 50 μL of a 1:10 dilution of the cleared lysate was denatured by adding 50 μL 2× sample buffer. Samples were analyzed by standard immunoblotting or frozen at −80°C.

### Plasmids, Molecular Cloning, Mutagenesis

For YFP-hMAD2, 600 ng total RNA of human primary mesenchymal stem cells were reversely transcribed using the RevertAid RT Reverse Transcription Kit (K1691, Thermo Scientific) according to the manufacturer. Subsequently, the sequence encoding for human MAD2 was amplified by PCR using primers flanked by restriction sites for BamHI and HindIII (BamHI_MAD2_rv: 5′- GTACGTGGATCCTCAGTCATTGACAGGAATTTTGTAGG-3′; HindIII_MAD2_fw: 5′- GTACGTAAGCTTATGGCGCTGCAGCTCTCCC-3′). PCR product (2 μg) and pEYFP-C1 vector DNA (2 μg) were digested with BamHI-HF and HindIII-HF (New England Biolabs) according to the manufacturer. Digested DNA was cleared from restriction enzymes using the NucleoSpin Gel and PCR Clean-up kit (740609.250, Macherey-Nagel). Insert-DNA (50 ng) was ligated with vector-DNA (150 ng) using the Fast-Link DNA Ligation Kit (LK0750H, Lucigen). XL10-Gold ultracompetent cells were transformed with the ligation product and streaked out on agar plates containing Kanamycin. Purified plasmid from resulting colonies was analyzed by control digestions with BamHI and HindIII and positive clones confirmed by sequencing (LGC Genomics).

For YFP-hGAT1, DNA was amplified by PCR using primers flanked by restriction sites for HindIII and KpnI (HindIII_hGAT1_fw: 5′- GACTGTAAGCTTTGGCGACCAACGGCAGCAAGGT-3′; KpnI_hGAT1_rv: 5′- GACTGTGGTACCCTAGATGTAGGCCTCCTTGCTGGTGG-3′). Subsequent steps were identical to YFP-hMAD2 cloning (see above) with KpnI-HF (New England Biolabs) replacing BamHI-HF. Digested PCR-product (150 ng) was ligated with digested pEYFP-C1 (150 ng).

For GST-hNET-Ct and GST-hDAT-Ct the sequences encoding the C-terminal 48 amino acids or 42 amino acids, respectively, were amplified from pEYFP-hNET and pEYFP-hDAT using primers encoding flanking restriction sites (EcoRI_hDATct_fw: 5′- ATATATGAATTCAAGTTCTGCAGCCTGCCTGG-3′; SalI_hDATct_rv: 5′- ATATATGTCGACCTACACCTTGAGCCAGTGGC-3′; BamHI_hNETct_fw: 5′- AGTCAAGGATCCAAAAGTTCCTCAGCACGCAGGG-3′; EcoRI_hNETct_rv: 5′- TGACAAGAATTCTCAGATGGCCAGCCAGTGTTG-3′). The PCR-product was inserted into the pGEX-5X-1 (GE Healthcare) according to the protocol above, using appropriate restriction enzymes (New England Biolabs). For both constructs 15 ng digested PCR-product were ligated with 150 ng digested vector. For all pGEX-5X-1 constructs, ampicillin was used for bacterial selection. Mutations were introduced by use of the QuikChange Lightning Site-Directed Mutagenesis Kit (Agilent Technologies) according to the manufacturer using the Agilent QuikChange Primer Design tool for primer design.

### Co-immunoprecipitation

HEK-293 cells stably expressing indicated YFP-tagged transporters were grown in 15 cm dishes to ~80% confluency. Dishes were placed on ice and washed once in ice-cold TBS. Ice-cold co-IP lysis buffer (500 μL) was applied, material scraped off the dish and transferred into 1.5 mL Eppendorf tubes. Cell lysis was completed at 4 °C under end-over-end rotation for 30 min. Lysates were cleared at 16 100 g at 4°C for 30 min. Meanwhile, GFP-Trap agarose (gta-100, Chromotek) (~ 12.5 μL packed resin per sample) was equilibrated in co-IP lysis buffer. Cleared lysates were incubate with GFP-Trap agarose for 90 min at 4 °C under end-over-end rotation. Agarose was washed thrice in 500 μL co-IP lysis buffer. Associated proteins were eluted in 50 μL 2× Laemmli SDS-PAGE sample buffer for 15 min at 60 °C. In the same step, 50 μL of a 1:10 dilution of the cleared lysate was denatured by adding 50 μL 2× sample buffer. Samples were analyzed by western blotting or frozen at −80 °C.

For co-IPs after siRNA-mediated MAD2-knockdown, 3 × 10^6^ HEK-293 cells stably expressing YFP-SERT in 10 cm dishes were transfected with siRNAs as outlined below. 2 days after transfection, cells were used for co-IP experiments as described above.

### siRNA transfection

HEK-293 cells in different culture formats were transfected with 3 target-specific siRNAs against human MAD2 (sc-35837, Santa Cruz Biotechnology) or siRNA against Luciferase (Dharmacon) as control. For 1 pmol of siRNA 0.3 μL of Lipofectamine RNAiMax transfection reagent (13778150, Invitrogen) were used for complex formation, according to the manufacturer. The final siRNA-concentration in the culture medium of each experiments was 10 nM. Gene silencing was allowed for 48 h. Subsequently, cells were subjected to further experimental procedures.

### Cell culture confocal microscopy

HEK-293 cells (1.25 × 10^4^ per well) stably expressing YFP-hSERT were seeded into 8-well μ-Slides (80826, ibidi). The following day, cells were transfected with siRNA as outlined above. Two days post siRNA-transfection, cells were imaged on a Zeiss LSM 510 laser scanning confocal microscope equipped with a 63× oil immersion objective, as described previously (Koban *et al.*, 2015). The relative intracellular YFP-SERT signal (RIS) was analyzed using ImageJ/polygonal selection software and calculated as:

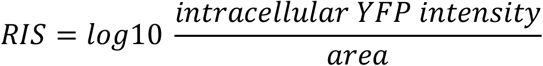

For co-localization with mCh-Rab5 and mCh-Rab7A, 1 day after siRNA transfection cells were transfected with according plasmids (25 ng per well) using 0.2 μL Lipofectamine 2000 transfection reagent (11668019, Invitrogen), according to the manufacturer. Cells were imaged on the following day. Co-localization was quantified using the JACoP-plugin for ImageJ (Bolte & Cordelieres, 2006) for calculating the Manders’ co-localization coefficient (Manders *et al*, 1993) corresponding to the fraction of mCherry overlapping with YFP.

### Radioactive substrate uptake

YFP-SERT expressing HEK-293 cells (5 × 10^4^ pre well of a 48-well plate) were transfected with siRNA as outlined above. Two days after transfection, cells were washed in Krebs-HEPES buffer containing glucose (KHB; 10 mM HEPES, pH = 7.3; 120 mM NaCl; 3 mM KCl; 2 mM CaCl2; 2mM MgCl2; 2 mM glucose monohydrate). For non-specific uptake, cells were pre-incubated for 10 min in KHB containing the SERT-inhibitor paroxetine (10 μM). Subsequently, cells were incubated for 1 min with constant 0.2 μM radioactive [^3^H]5-HT in the presence or absence of non-tritiated (“cold”) 5-HT in order to reach total 5-HT-concentrations of 0.2, 1, 3, 10 and 30 μM. Non-specific uptake was determined with the 30 μM concentration in the presence of 10 μM paroxetine. Wells were washed with 500 μL ice-cold KHB, cells were lysed in 1% SDS and radioactivity was measured by liquid scintillation counting. Intact cells on the same well-plate, seeded and transfected in the same manner, were trypsinized and counted using a hemocytometer. Specific [^3^H]5-HT uptake was calculated as pmol/10^6^ cells/min. K_M_ and V_max_ were determined by non-linear regression according to Michaelis-Menten kinetics. Residual trypsinized cells were lysed after trypsin-inactivation and used to determine the efficiency of siRNA-mediated knockdown.

### Surface biotinylation and reversible biotinylation

#### Surface biotinylation

YFP-SERT expressing HEK-293 cells (6 × 10^5^ per well) in 6 cm dishes were transfected with siRNA as outlined above. After 2 days, cells were washed in PBS additionally containing 1 mM MgCl_2_ and 0.1 mM CaCl_2_ (PBS^2+^). Washed cells were incubated with PBS^2+^ containing 2 mg mL^−1^ Pierce Premium Grade Sulfo-NHS-SS-Biotin (PG82077; Thermo Scientific) for 30 min on ice. Reaction was quenched twice for 15 min with 100 mM glycin in PBS^2+^ and cells were washed thrice in TBS. Cells were lysed in 300 μL RIPA-buffer (20 mM Tris-HCl, pH = 7.6; 150 mM NaCl; 1 mM EDTA; 1% Triton X-100; 0.1% SDS; 0.5% sodium deoxycholate; protease inhibitor cocktail; phosSTOP) per well. Lysates were cleared at 16 100 g for 30 min at 4°C. Meanwhile, high capacity streptavidin agarose (20357, Thermo Scientific) (~ 25 μL packed resin per sample) was equilibrated in RIPA-buffer. 30 μL cleared lysate was saved as separate sample and residual lysate was incubated with streptavidin agarose overnight at 4°C under end-over-end rotation. Agarose was washed thrice in 500 μL RIPA-buffer lacking sodium deoxycholate and protein/phosphatase inhibitors. Immobilized biotinylated proteins were eluted in 100 μL 2× Laemmli SDS-PAGE sample buffer at 90°C for 10 minutes. In the same step the 30 μL lysate sample was denatured by adding 30 μL 2× sample buffer. Samples were either analyzed directly by western blotting or stored at −80°C.

#### Reversible biotinylation

YFP-DAT expressing HEK-293 cells were siRNA-transfected and biotinylated as for surface biotinylation. After quenching in 100 mM glycine, one 6-well plate containing the “total”- and the “strip”-controls remained on ice. The other 6-well plate containing the “internalization”-samples was moved to room temperature and washed once with 37°C warm internalization solution (PBS^2+^ supplemented with 0.2% (w/v) IgG and protease free bovine serum albumin [A3059, Sigma-Aldrich], 0.18% (w/v) glucose monohydrate, 500 nM phorbol 12-myristate 13-acetate). Internalization solution (1 mL) was applied to each well and proteins were allowed to internalize at 37°C 5% CO_2_ for 15 min. “Internalization”-samples were put back on ice and all wells were washed in ice-cold NT-buffer (20 mM Tris-HCl, pH = 8.0; 150 mM NaCl; 1 mM EDTA). The cellular surface was stripped from the biotinylation (except for the “total”-controls) in NT-buffer containing 150 mM Tris(2-carboxyethyl)phosphine (pH = 8.1) two times for 15 min. Samples were washed extensively in ice-cold TBS and cells were lysed in 700 μL RIPA-buffer. Afterwards, biotinylated proteins were pulled down as for surface biotinylation. Immunoblots were analyzed using ImageJ and internalization rates (IR) were calculated as:

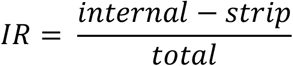

### Statistical analysis

As appropriate, unpaired or paired two-tailed Student’s t-tests or one-way ANOVA with Dunnett’s multiple posthoc comparison or Pearson’s correlations were conducted using GraphPad software. P-values below 0.05 were considered significant. The specific statistical tests as well as the exact p-values are indicated in the figure legends.

## Acknowledgements

We express our gratitude to Hongtao Yu and Eunhee Choi for providing antibodies against human and mouse p31^COMET^. We thank Kathrin Jäntsch and Melanie Burger for assistance with radioactive substrate uptake and molecular cloning of YFP-GAT1, respectively. We acknowledge financial support from the Austrian Science Fund FWF project P31255-B27 to Sonja Sucic for providing the DNA for human GAT1. This work was funded by the Vienna Science and Technology Fund (WWTF) project LSC17026.

## FIGURE LEGENDS

**Fig. EV1.**
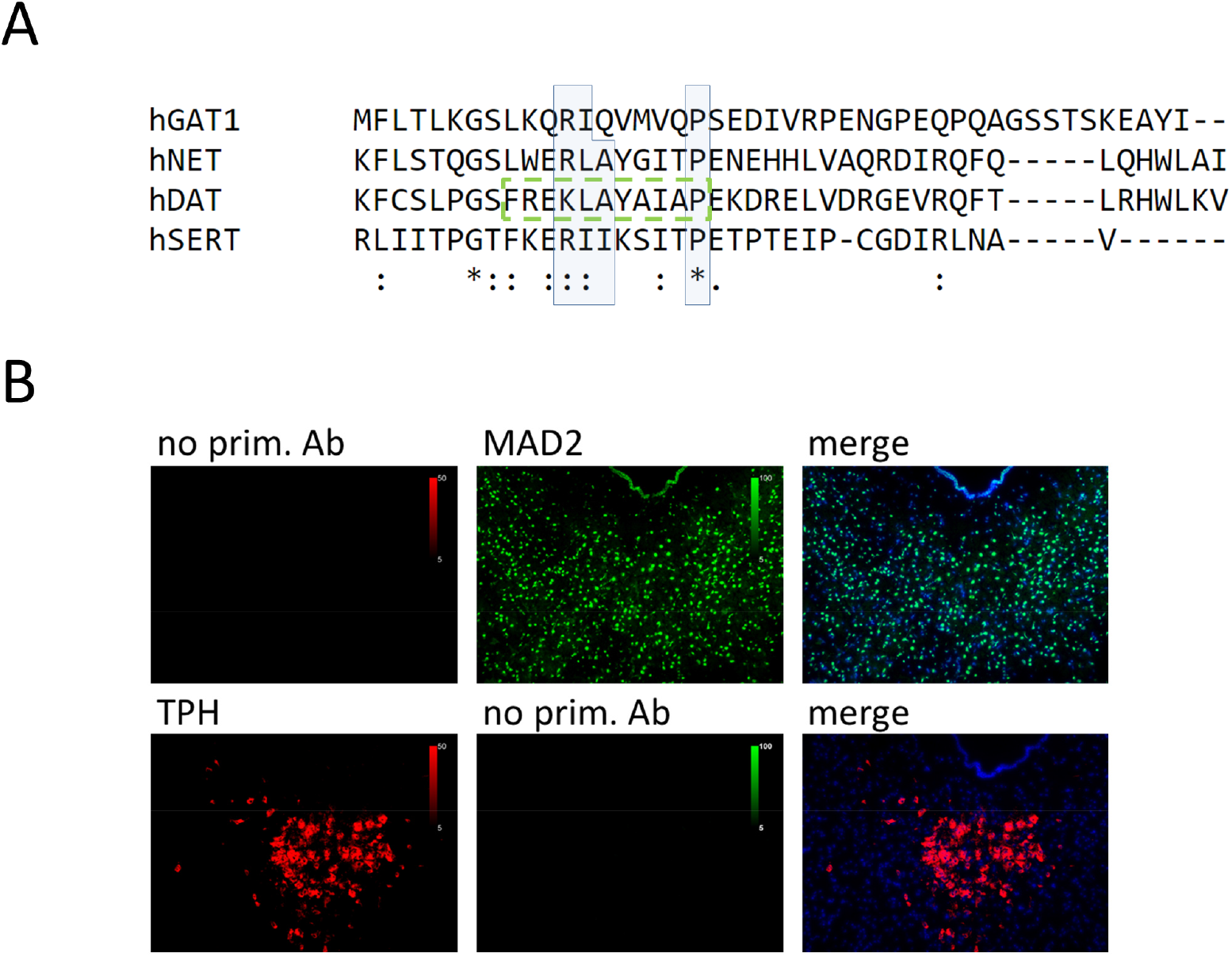
Conservation of transporter C-terminal MIMs. “No primary antibody”-controls. A Full length C-termini of indicated transporters were aligned using Clustal Omega software. Conserved residues of possible MIMs are highlighted by blue boxes; (**\***) - fully conserved residue; (:) - residues of strongly similar properties; (.) residues of weakly similar properties. The previously described endocytic motif in DAT is highlighted by the green dashed box. B Mouse brain sections were subjected to immunofluorescence microscopy as described in “*Materials and Methods*”, omitting polyclonal antibodies for either TPH or MAD2.

**Fig. EV2.**
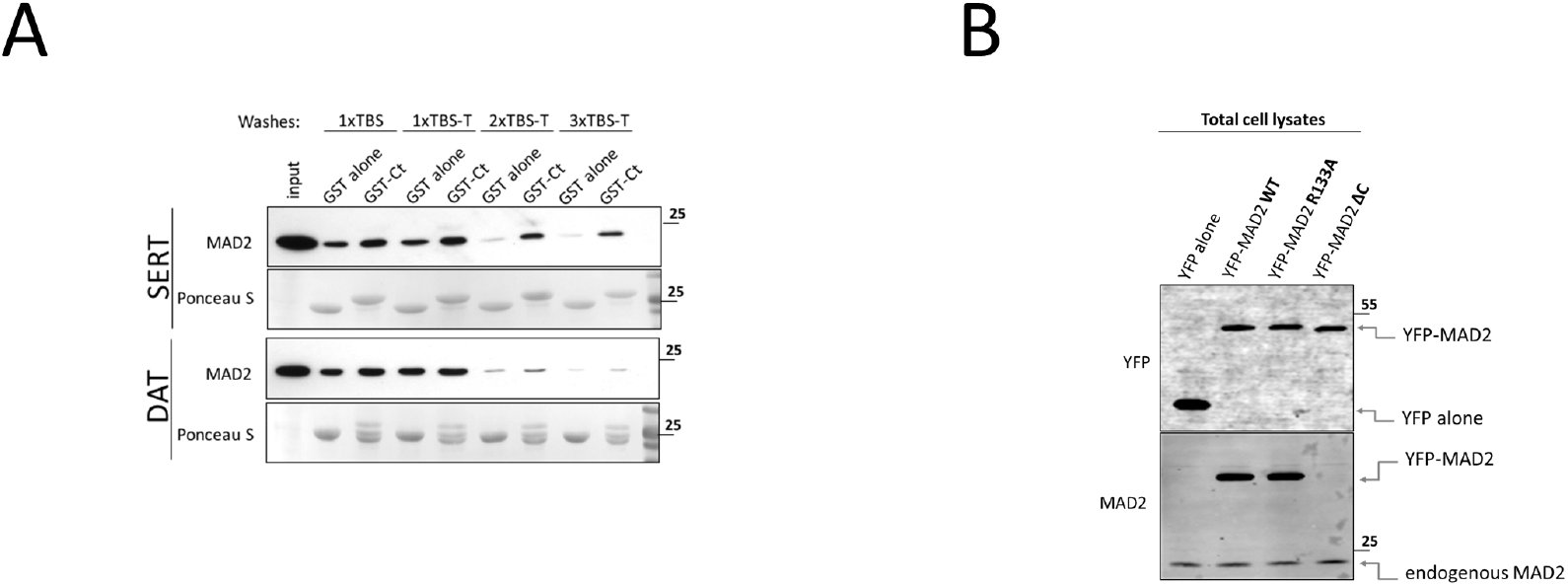
MAD2-association with SERT- and DAT-C-termini. MAD2 C-terminal deletion disrupts antibody recognition. A GST pull-down experiments were conducted as described in “*Materials and Methods*”. Following incubation with HEK-293 lysate, reactions were washed up to three times in the indicated buffer systems. Bound proteins were eluted and analyzed by immunoblotting. B Total cell lysates (5 μg) of indicated transfections were subjected to immunoblotting, using antibodies against YFP and MAD2.

**Fig. EV3.**
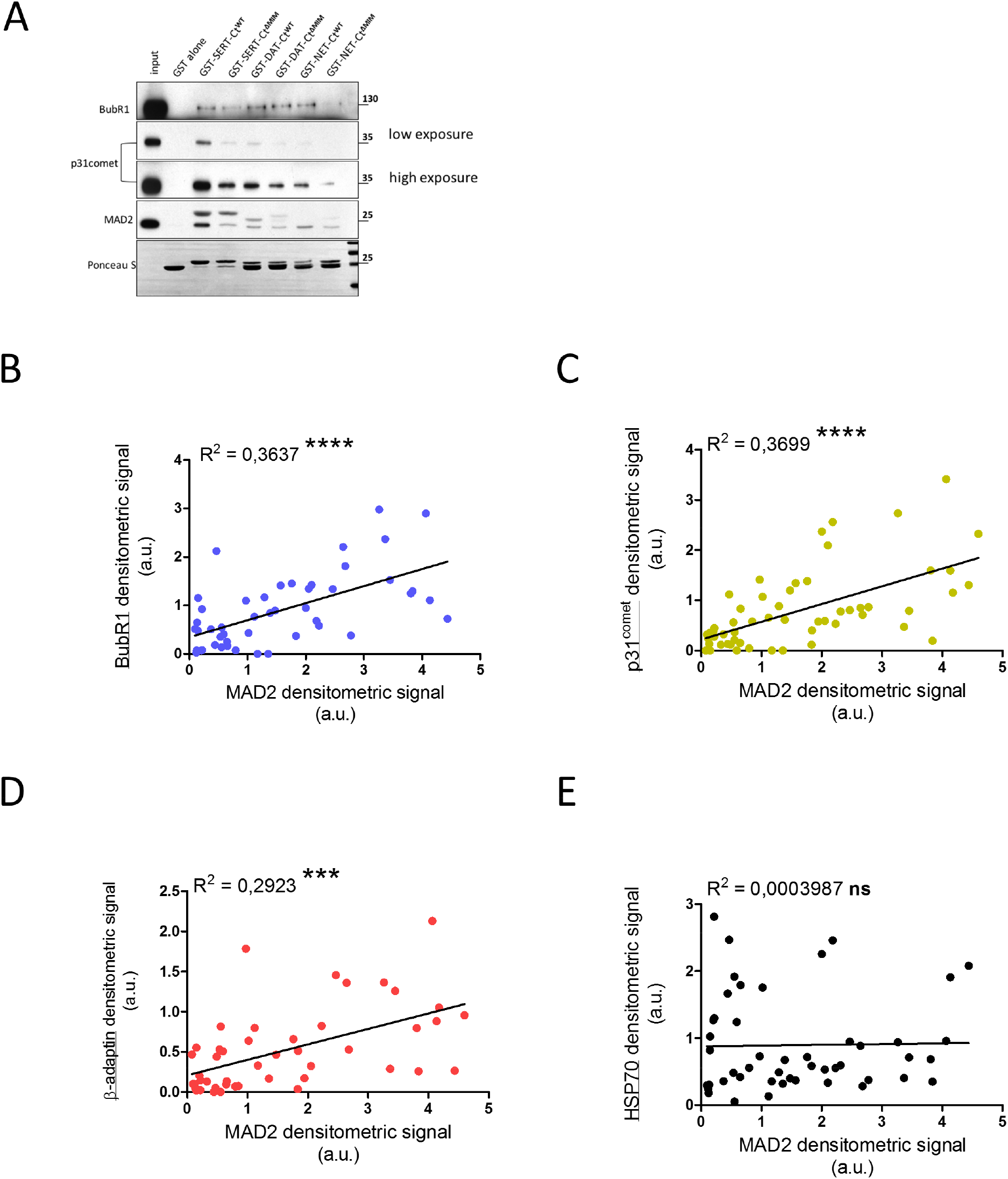
Association of full-length transporters with BubR1, β2-adaptin and p31^COMET^ correlates with MAD2-association. A GST pull-down experiments were conducted and the levels of BubR1 and p31^COMET^ in eluates were analyzed by immunoblotting. The range between the highest (GST-SERT-Ct^WT^) and the lowest (GST-NET-Ct^ΔMIM^) amount of pulled down p31^COMET^ prompted us to include two different exposures of the same immunoblot. B – D Scatter plots for the co-IP of indicated proteins (y-axis) against the co-IP of MAD2 (x-axis) (cf. Fig. 3 E – J). R^2^ indicates squared Pearson’s correlation coefficients; ***p < 0.001, ****p < 0.0001, ns – not significant.

**Fig. EV4.**
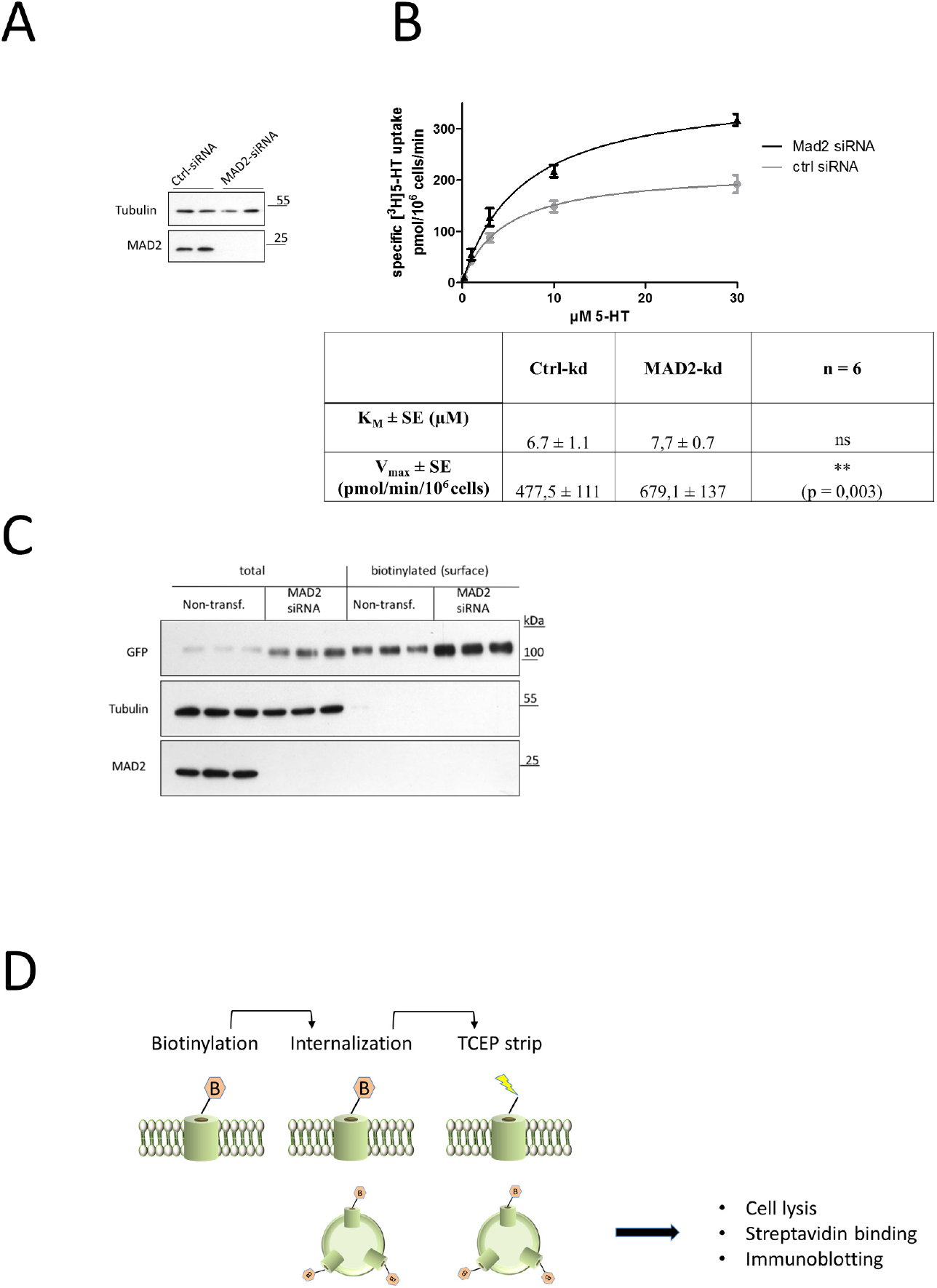
MAD2-kd increases SERT surface levels but does not affect transport-activity. A - B Radioactive substrate uptake was conducted as described in “*Materials and Methods*”. K_M_ and V_max_ were determined by fitting the data to the equation of a rectangular hyperbola. A representative graph is shown in (B). Paired two-tailed t-test; **p < 0.01, ns – not significant; exact p-value is indicated. C Cell surface biotinylation using HEK-293 cells stably expressing YFP-hSERT was performed as described in “*Materials and Methods*”. Whole cell lysates (n = 11) and streptavidin-conjugated fractions (i. e. surface protein) (n = 4) were subjected to immunoblotting. D Schematic sequence of reversible biotinylation experiments, as conducted for Fig. 5 E.

## Notes

### Competing Interest Statement

The authors have declared no competing interest.

